# Tetrodotoxin-resistant Na_V_1.8 contributes to action potential conduction in a population of mouse and human vagal nerve fibers

**DOI:** 10.64898/2026.11.19.767887

**Authors:** Sanjay S. Nair, Thomas E. Taylor-Clark

## Abstract

Axonal action potential (AP) conduction is dependent on voltage-gated sodium channels (Na_V_1s). AP conduction in the vagus nerve (which contains afferent and efferent fibers) is essential to its reflex regulation of most visceral organs. The nine isoforms of Na_V_1s show diverse expression patterns in vagal neurons. In rodent vagus nerves, multiple Na_V_1 isoforms contribute to AP conduction, with Na_V_1.7 playing an essential role in ∼70% of fibers. Here, we investigated the contribution of tetrodotoxin-resistant (TTX-R) Na_V_1.8 towards vagal AP conduction ex vivo. We tested two structurally different Na_V_1.8-selective inhibitors, A803467 and PF01247324 and their effects on electrically evoked C-fiber compound action potentials (CAPs) along mouse and human vagus nerves. Tetrodotoxin completely abolished mouse and human vagal CAPs. Nevertheless, both A803467 and PF01247324 caused a concentration-dependent decrease in the C-fiber CAPs in mouse and human vagus nerves. The selectivity of both drugs was tested in Na_V_1.8 knockout mice. A803467 but not PF01247324 significantly decreased Na_V_1.8 KO vagal CAPs, indicating that PF01247324’s inhibition of wildtype vagal CAPs was solely mediated by Na_V_1.8, and that A803467 caused inhibition via Na_V_1.8-mediated and non-Na_V_1.8-mediated mechanisms. In further studies of wildtype mice, Na_V_1.8 inhibition following incubation with Na_V_1.7-selective inhibitor PF-05089771 showed no further decrease in mouse vagal CAP, demonstrating that Na_V_1.8-dependent inhibition affects fibers also sensitive to Na_V_1.7-dependent inhibition. These data suggest that TTX-sensitive Na_V_1.7 and TTX-R Na_V_1.8 cooperate to conduct APs along a select population of vagal nerves. Axonal Na_V_1.8 inhibition could be utilized to treat aberrant vagal nerve activity in visceral organ diseases.

## Introduction

The generation and conduction of neuronal action potentials (APs) is critically dependent on voltage-gated sodium channels (Na_V_1s). There are nine pore-forming alpha Na_V_1 subunits in mammals, ranging from Na_V_1.1 to Na_V_1.9, which show varying levels of expression in neurons (Catterall et al., 2005). These Na_V_1s are classified by their sensitivity to pufferfish toxin tetrodotoxin (TTX) inhibition. TTX-resistant (TTX-R) Na_V_1s include Na_V_1.5, Na_V_1.8 and Na_V_1.9, while the remaining TTX-sensitive (TTX-S) Na_V_1s are inhibited by nanomolar concentrations of TTX. The vagus nerve, being a mixed peripheral nerve, consists of both afferent (80%) and efferent fibers (20%) that regulate the functions of many visceral organs. These constituent nerve fibers can be divided into two groups based on their physical structure: the myelinated, fast conducting (>1.5m/s), large-diameter A-fibers and the unmyelinated, slow-conducting (<1.5m/s), small-diameter C-fibers. In the vagus, C-fibers outnumber A-fibers by a ratio of ∼8:1 (Mazzone & Undem, 2016). Vagal sensory afferents have their neuronal soma in one of two sensory ganglia (nodose ganglion or jugular ganglion). In general, A-fiber vagal afferents are thought to control homeostatic, physiological functions, whereas C-fiber vagal afferents have nociceptive characteristics, mediating defensive reflexes. Excessive or aberrant vagal nerve activity, especially in nociceptive vagal C-fibers, has been shown to contribute to many diseases affecting vagally-innervated visceral organs such as asthma, chronic cough and gastrointestinal dysmotility (Mazzone & Undem, 2016; Prescott & Liberles, 2022). Vagal Na_V_1s are attractive therapeutic candidates as they are critical for neural activity. Understanding the role of specific Na_V_1 isoforms in AP initiation and conduction in vagal nerves is important for two reasons: firstly, it provides the mechanistic rationale for targeting dysfunctional vagal nerves in diseased states; secondly, it provides a rationale for avoiding side effects associated with targeting specific Na_V_1s for the treatment of pain and CNS disorders such as epilepsy (Alsaloum et al., 2020, 2025).

Initial electrophysiological studies of the vagus nerve demonstrated the complete abolition of AP conduction by TTX, with isoform selective inhibitors suggesting a major role for Na_V_1.7 (Hadley et al., 2021; Kollarik et al., 2022; Muroi et al., 2011; Ru et al., 2020). These studies were interpreted by many to indicate that (1) TTX-S Na_V_1 mediated AP conduction and (2) TTX-R Na_V_1 were not required for AP conduction. Nevertheless, Na_V_1.8 is expressed in almost all vagal neurons (Gautron et al., 2011; Kwong et al., 2008; Muroi et al., 2011) and patch clamp studies of nodose vagal neurons show that Na_V_1.8-like Na+ currents are capable of evoking APs in the presence of TTX (Kwong et al., 2008). Recently, we confirmed the abolition of AP conduction by TTX in mice and rat vagus nerves (Nair et al., 2023). Importantly, this study also showed that A-803467 (A80), a drug shown to be >100-fold selective for human Na_V_1.8 over TTX-S Na_V_1s in patch clamp studies (Jarvis et al., 2007), blocked vagal AP conduction in ∼40% of vagal axons (Nair et al., 2023). This suggested that in some vagal fibers both TTX-S channels and Na_V_1.8 were required for AP conduction. Here, we further investigated the role of axonal Na_V_1.8 in AP conduction in mouse and human vagal C-fibers. For this we used two structurally different Na_V_1.8-selective inhibitors A-803467 (A80) and PF-01247324 (PF24), in combination with a Na_V_1.8 knockout mouse model. Our data suggests that selective inhibition of Na_V_1.8 blocks AP conduction in a subset of vagal nerve fibers. We also show that Na_V_1.8-dependent block affects vagal C-fibers that are sensitive to Na_V_1.7 inhibition.

## Methods

All animal experiments were performed with approval from The University of South Florida Institutional Animal Care and Use Committee (AAALAC No. 000434). 6-8 weeks old male wild-type C57BL/6J mice (from Jackson Laboratories, #000664) and homozygous Na_V_1.8 Cre knock-in/knock-out mouse *B6.129(Cg)-Scn10atm2(cre)Jwo/TjpJ* (Na_V_1.8 KO, from Jackson Laboratories, #036564) were used for our experiments. Na_V_1.8 KO mice carry a Cre recombinase cassette that was inserted into the start site of the Scn10a gene’s translation locus (Nassar et al., 2004). This insertion replaces the methionine start codon thereby abolishing Na_V_1.8 expression. Mice were housed in their respective cages under normal laboratory conditions of 20C, on a 12-hour light/dark cycle. Mice were given standard rodent chow and water ad libitum.

Mice were euthanized by CO2 inhalation and exsanguination. The cervical and thoracic vagus nerve was isolated, then 7-10mm in length was dissected out, starting immediately distal to the nodose vagal ganglion. The vagus nerve from either side were used for our experiments. Human vagus nerves samples were obtained from deidentified organ donor lungs (donor information in Table X) in collaboration with the Tampa-based nonprofit LifeLink organization, which operates federally certified organizations to recover human organs for transplantation. Lungs (with attached bronchi) were extracted by certified surgeons according to the standard transplant protocols and transported in SPS1 solution, also known as UW solution. Samples were exclusively used within a 12-hour window following organ-retrieval surgery. For the human vagus, nerves were taken from the thoracic level and carefully separated into individual fascicles for our recordings. Some experiments also used bronchial branches of the vagus.

For both mouse and human nerves, the epineural sheath was removed carefully under the microscope using bent forceps to allow complete access for the drugs to its intended targets. Care was taken not to stretch/damage the nerve fibers during the sheath removal process. The dissected vagi were superfused with Krebs buffer solution (KBS; in mM: 118 NaCl, 5.4 KCl, 1 NaH2PO4, 1.2 MgSO4, 1.9 CaCl2, 25 NaHCO3, and 11 Dextrose), gassed with 95% O2 and 5% CO2 in a Sylgard-lined, water-jacketed (maintained at 35oC) organ bath. The buffer solution was also supplemented with 3uM indomethacin to minimize the influence of autacoids released during dissection. The distal end of the nerve was pinned down to the Sylgard using a fine dissection pin. The proximal end of the vagus nerve was placed in a suction recording electrode to record compound action potentials (CAPs). The recording electrode bore was sealed with Vaseline to form a high-resistance electrical seal, greatly improving the signal to noise ratio of our CAP recordings. The suction electrode consisted of a silver (Ag) recording wire attached to a glass microelectrode preamplifier holder. An AC amplifier (A-M Systems Microelectrode AC Amplifier, MODEL 1800) connected to the microelectrode holder, amplified and filtered (<1 Hz, >1 kHz) the signal. The recorded signal was then processed and visualized using an oscilloscope (Tektronix, TDS 3054B). An A-M Systems isolated pulse stimulator (MODEL 2100) delivered a square pulse of 0.2 ms duration to the nerves. Vagus nerves were stimulated with voltages ranging from 30-60V using a concentric electrode made of platinum and iridium (PI-NE-100, MicroProbes, Gaithersburg, MD). Once a certain voltage was used to obtain control CAP recordings, it was left unchanged for the remaining duration of the experiment. The entire circuit was grounded using a silver-silver chloride (Ag-AgCl) pellet that was placed in the perfusion bath next to the recording electrode.

A803467 (A80) stock solution was made up in 100% ethanol. PF-05089771 (PF71) and PF-01247324 (PF24) stock solution was made up in DMSO. Final working concentrations were obtained by diluting them in KBS on the day of the experiments. We have chosen to use concentrations greater than the IC50 published in the original patch clamp characterization of these selective Na_V_1 inhibitors. All the inhibitors used in this study apart from TTX are state-dependent inhibitors of their respective Na_V_1 (Alexandrou et al., 2016; Jarvis et al., 2007; Payne et al., 2015). IC50s that are available in the literature are determined from patch-clamp studies of native and recombinant Na_V_1s where these channels can be driven into a preferred state (usually the inactive state). It is not possible to know the biophysical state of the Na_V_1s in our ex vivo preparations. Moreover, higher concentrations are needed in these ex-vivo studies to overcome diffusion barriers in tissues.

### Stimulation Protocol

The CAP was recorded during 15-s periods of 1 Hz; at all other times the nerve was only electrically stimulated once per minute (0.0166 Hz).

The general experimental protocol was as follows: vagal incubation with KBS for 15 min, followed by 1st control stimulation period (C1), then another 15 min of KBS incubation, followed by the 2nd control stimulation period (C2). After two consecutive reproducible control CAPs were recorded, the vagus was incubated for 30 min with the Na_V_1.8 inhibitor or Na_V_1.7 inhibitor or DMSO, and another CAP was recorded. For concentration-response curves and inhibitor combination experiments, a higher concentration of the respective inhibitor or a second inhibitor was applied for 30mins, which was followed by another CAP recording. Finally, all protocols ended as follows: 30min incubation of 300nM TTX, a CAP recording, then a nerve pinch between the stimulating and recording electrodes to destroy all conduction, followed by a final CAP recording.

### Data Analysis

The CAP signals were recorded and analyzed by Spike 2 software (Cambridge Electronic Design, England). The length of the nerve between the stimulating and recording electrodes was recorded for each experiment to determine the conduction velocity of the CAP components. We have opted to analyze only the C-fiber CAP component in this study. Any part of the CAP that was <1.5m/s was categorized as the C-wave component of the vagal CAP. The positive and negative deflections of the C-wave CAP were transformed by squaring and square rooting the amplitude against time waves of three consecutive CAPs generated during the 15-second, 1Hz stimulation train. The “transformed” CAPs were then averaged and the area under the curve (AUC) for the C-waves were quantified. The AUCs calculated after the nerve pinch were subtracted from the AUCs determined for each test [i.e., control 1 (C1), control 2 (C2), Na_V_1 inhibitors, and TTX]. Pinch-adjusted AUCs were then expressed as a percentage of the control C1 CAP. Individual data points were only included if the C2 AUC C-waves were reproducible (compared with C1). All the data are presented as mean ± SD.

GraphPad Prism and Microsoft Excel were used for statistical analysis. In most cases, paired groups were compared, with one-way repeated-measures ANOVA with Sidak’s multiple comparisons test. Post hoc comparisons to the control C2 group after ANOVA was conducted using Dunnett’s post-hoc test. Groups from separate experiments were compared with a 2-tailed unpaired t-test with Welch’s correction. A P value less than 0.05 was considered significant.

## Results

### Action potential conduction in mouse vagal C-fibers

The conduction of electrically evoked action potentials along the mouse vagus nerve was determined in ex vivo compound action potential (CAP) recordings. CAP signals were stable over 2-hour experiments. Incubation with 300nM TTX completely abolished the mouse C-wave CAP signal. Partial recovery of the C-wave CAP was observed after wash-out of TTX block. The residual CAP signal after treatment with TTX (prior to washout) was not significantly different from the CAP signal post-nerve pinch (TTX: -0.24±1.6% of C1 vs Pinch: 0% of C1, Fig.1, n=4, P=0.99), indicating that TTX-S Na_V_1s are essential for all C-fiber AP conduction in the mouse vagus nerve.

**Figure 1:**
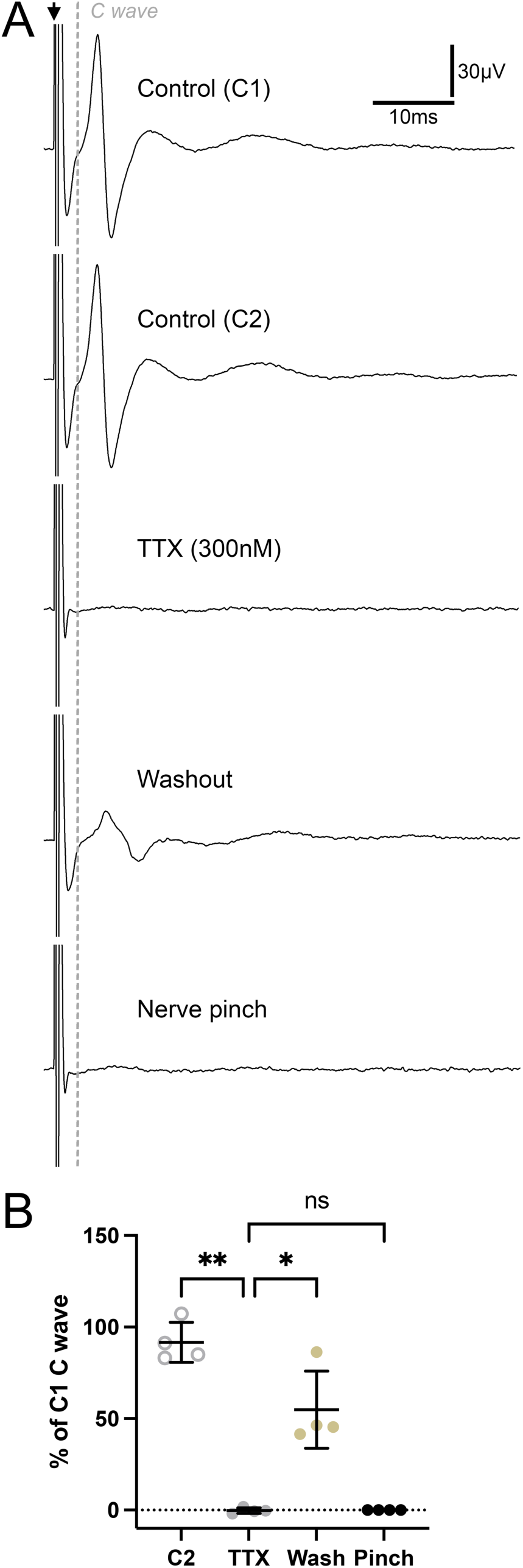
Effect of TTX on mouse vagal compound action potential (CAP). A, representative mouse vagal electrically evoked CAP during 1^st^ control (C1), 2^nd^ control (C2), after 30min 300nM TTX, after 30min washout and after destruction of the nerve. Black arrow denotes the shock artifact from electrical stimulation. Vertical grey line represents the division of A- and C-wave based on the length of the nerve between the two electrodes (<1.5m/s considered as C-wave). B, mean ± SD of the AUC of the C-wave CAP normalized to the initial control (C1) during C2 control, after TTX (300nM), after washout with Krebs, and after nerve pinch (n=4). One-way repeated-measures ANOVA with Sidak’s multiple comparisons test *P<0.05, **P<0.005, ns P>0.05.

We repeated our previous study (Nair et al., 2023), again showing that 1µM of the Na_V_1.8 inhibitor A80 caused ∼35% reduction of the vagal C-wave CAP (C2: 99.0±13.0% of C1 vs A80 1µM: 64.5±22.2% of C1, Supplemental Fig. 1, n=15, P=0.0004). We then showed that 30min incubation of A80 blocked the C-fiber component of the CAP in a concentration-dependent manner (Fig. 2A & 2B). While 0.3µM A80 had minimal effect (p=0.095), 1µM A80 significantly reduced the C-wave CAP signal by ∼30% (C2: 99.7±9.7% of C1 vs A80 1µM: 73.2±18.2% of C1, n=7, P=0.028) and 3µM concentration caused a 50% reduction of the C-wave CAP when compared to C2 control CAP (C2: 99.7±9.7% of C1 vs A80 3µM: 50.1±21.9, n=7, P=0.0074). The residual C-wave CAP was abolished by subsequent TTX incubation (P<0.0001). Altogether, this data suggests that mouse Na_V_1.8 is required for AP conduction in a subset of vagal C-fibers.

**Figure 2:**
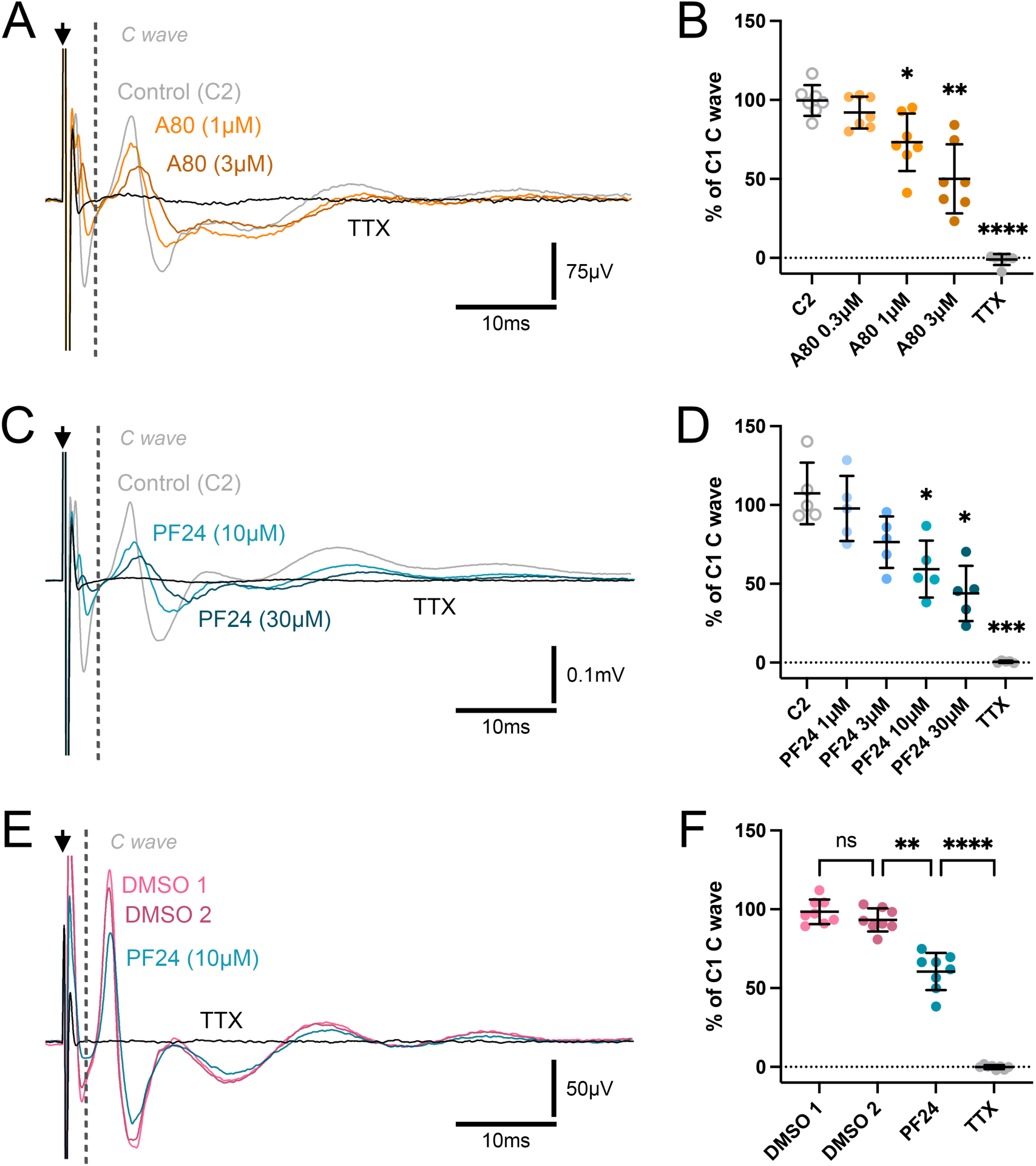
Effect of selective Na_V_1.8 inhibitors on mouse vagal compound action potential (CAP). A, representative mouse vagal CAP during control C2 (grey trace), after 30min treatment with escalating doses of A80 (orange and brown traces) and after 300nM TTX (black trace). B, mean ± SD of the AUC of the C-wave CAP normalized to the initial control (C1) during C2 control, A80 (0.3 to 3µM) and TTX (n=7). C, representative mouse vagal CAP during control C2 (grey trace), after 30min treatment with escalating doses of PF24 (teal and blue traces), and after 300nM TTX (black trace). D, mean ± SD of the AUC of the C-wave CAP normalized to the initial control (C1) during C2 control, PF24 (1 to 30µM) and TTX (n=5). E. Representative mouse vagal CAP after 1^st^ DMSO treatment, 2^nd^ DMSO treatment, 10µM PF24, and 300nM TTX. F, mean ± SD of the AUC of the C-wave CAP normalized to the initial control (C1) during sequential DMSO administrations, 10µM PF24 and TTX (n=8). In A, C and E, black arrow denotes the shock artifact; vertical grey line represents the division of A- and C-wave. In B, D and F, one-way repeated-measures ANOVA with Sidak’s multiple comparisons test, compared to respective C2 control values *P<0.05, **P<0.005, ****P<0.0001, ns P>0.05.

We then tested a structurally different Na_V_1.8 inhibitor, PF24, on AP conduction in mouse vagus nerves (Fig. 2C & 2D). While 30min incubation with 1µM and 3µM PF24 had little effect (P=0.083 and 0.096, respectively), 10µM PF24 caused ∼40% reduction in mouse C-wave CAP (C2: 107.4±19.6% of C1 vs PF24 10µM: 59.3±18.1% of C1n=5, P=0.031). Although 30µM PF24 inhibited the C-wave CAP signal by ∼55% in comparison to the control C-wave CAP (C2: 107.4±19.6% of C1 vs PF24 30µM: 43.9±17.5% of C1, n=5, P=0.016), it did not abolish it. The residual CAP signal was abolished by 300nM TTX (P=0.0008). These findings are consistent with the A80 data, further supporting the conclusion that Na_V_1.8 is required in the conduction of APs in a population of murine vagal C-fibers.

We also tested the effect of consecutive DMSO incubations (the vehicle for Na_V_1.8 inhibitors) to check for the possibility of non-specific CAP signal degradation in our ex-vivo preparations. Incubation with 0.2% DMSO for 30mins and 1hr had no effect on the control C-wave CAP recordings (Fig. 2E & 2F, n=8, P=0.29). Subsequent incubation with 10µM PF24 inhibited the C-wave CAP by ∼40% (2^nd^ DMSO: 93.3±7.3% of C1 vs PF24 10µM: 60.5±11.8% of C1, n=8, P=0.002). The residual CAP signal was once again abolished by subsequent TTX treatment (P<0.0001).

To determine the selectivity of the Na_V_1.8 inhibitors A80 and PF24 employed in our study, we tested these two inhibitors on electrically evoked CAPs recorded from vagus nerves taken from Na_V_1.8 KO mice. Electrical stimulation of these vagus nerves produced reproducible C-wave CAPs. Incubation with 1µM A80 for 30mins significantly inhibited ∼20% of the Na_V_1.8 KO C-wave CAP (C2: 99.2±3.5% of C1 vs A80 1µM: 83.2±12.9% of C1, Fig. 3A & 3B, n=11, P=0.0064). The residual CAP signal was abolished by TTX (P<0.0001). We compared the A80-mediated inhibition of vagal C-wave CAP between WT and Na_V_1.8 KO experiments using an unpaired t-test of the aggregate data. A80-mediated inhibition was significantly reduced in Na_V_1.8 KO vagus nerves (67.2±21.0% of C1 in WT (n=21) vs 83.2±12.9% of C1 in Na_V_1.8KO mice (n=11), Supplemental Fig. 2, P=0.011). Thus, the reduction of vagal C-wave CAP in wild-type vagi by 1µM A80 is via an Na_V_1.8-mediated mechanism and a non-NaV1.8-mediated mechanism. Interestingly, incubation of 10µM PF24 caused no reduction in the Na_V_1.8 KO C-wave (C2: 100.2±2.0% of C1 vs PF24 10µM: 102.7±13.9% of C1, Fig. 3C & 3D, n=6, P=0.95). Thus, unlike A80, PF24’s reduction of the wild-type vagal C-wave CAP occurs only via the inhibition of Na_V_1.8.

**Figure 3:**
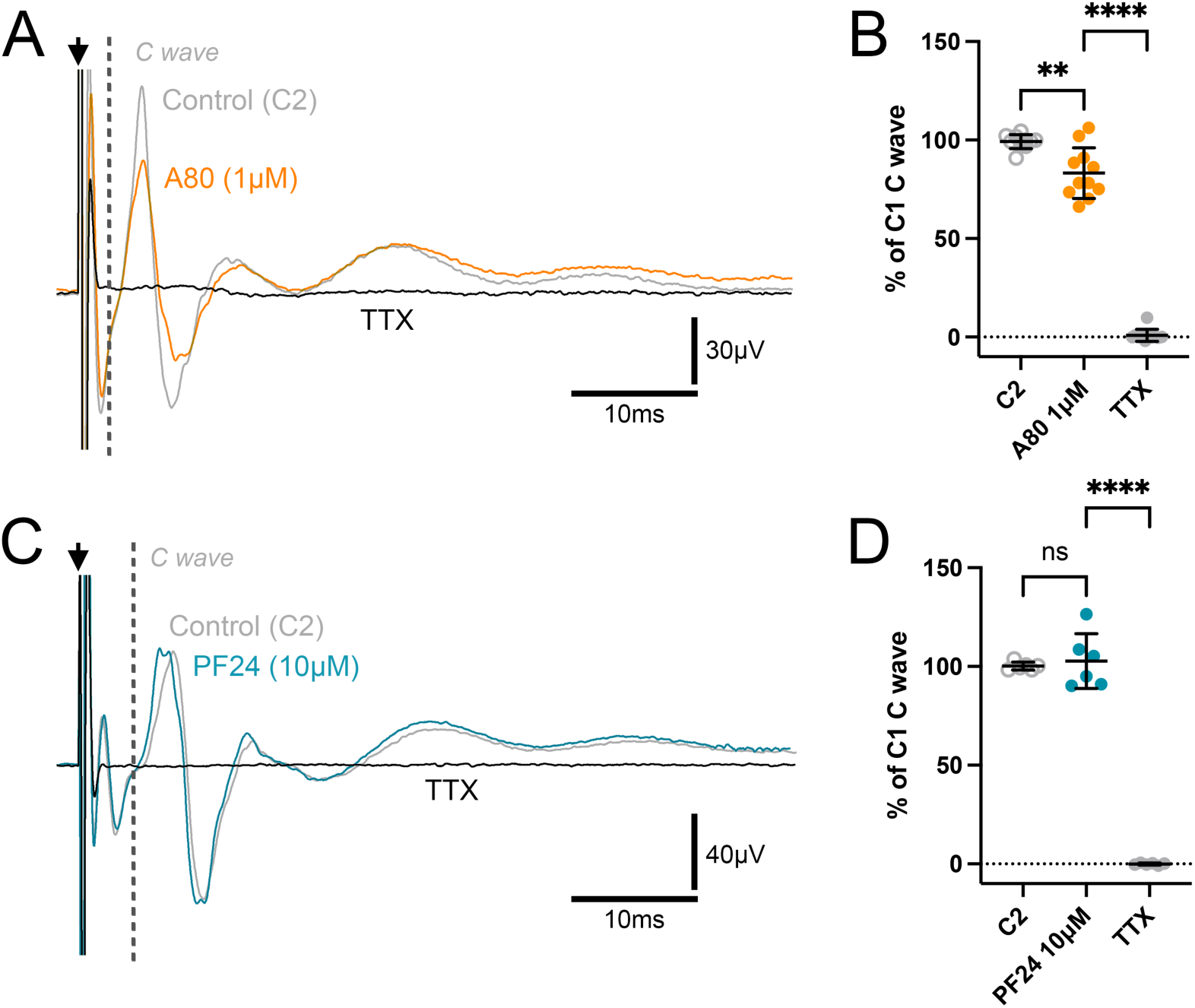
Effect of A80 and PF24 on Na_V_1.8 KO mouse vagal compound action potential (CAP). A and C: Representative Na_V_1.8 KO mouse vagal CAP during control C2 (grey trace), after 30min treatment with 1µM A80 (A, orange trace) or 10µM PF24 (C, teal trace), and after 300nM TTX (black trace). Black arrow denotes the shock artifact; vertical grey line represents the division of A- and C-wave. B and D, mean ± SD of the AUC of the C-wave CAP normalized to the initial control (C1) during C2 control, 1µM A80 (B, n=11) or 10µM PF24 (D, n=6) and TTX. One-way repeated-measures ANOVA with Sidak’s multiple comparisons test **P<0.005, ****P<0.0001, ns P>0.05.

We have previously shown that Na_V_1.7 is a key TTX-S contributor to AP conduction along rodent vagus nerves (Nair et al., 2023). To study the phenomenon of cooperativity between TTX-S Na_V_1.7 and TTX-R Na_V_1.8 in C-fiber AP conduction, we tested a combination strategy of treating WT mouse vagi with the Na_V_1.7 selective inhibitor PF71 followed by the Na_V_1.8 inhibitors tested above. 30min incubation with 3µM PF71 caused a significant reduction (∼65%, P<0.0001) in the C-wave CAP, similar to previous studies. Subsequent addition of 10µM PF24 to PF71-treated vagi failed to cause further significant reductions in the C wave (PF71 3µM: 37.2±12.3% of C1 vs PF71 + PF24 10µM: 25.5±14.1% of C1, Fig 4A & 4B, n=6, P=0.12). Subsequent TTX incubation abolished the C wave CAP (n=6, P=0.026). The lack of effect of PF24 in combination with PF71 suggests that vagal nerve fibers that are sensitive to Na_V_1.8-dependent inhibition are a subset of fibers that are also dependent on Na_V_1.7. In another set of experiments, addition of 1µM A80 in combination with PF71 caused a significant reduction in the C-wave compared to that caused by PF71 alone (PF71 3µM: 42.1±12.1% of C1 vs PF71 + A80 1µM: 13.8±6.7% of C1, n=5, Fig. 4C & 4D, P=0.012). Subsequent TTX incubation caused further significant reduction and abolished the C wave (P=0.014). These data are consistent with the Na_V_1.8 KO data that suggests A80 reduces the C wave via at least one non-Na_V_1.8-mediated mechanism. Furthermore, these data suggest that the non-Na_V_1.8-mediated mechanism of A80 in part reduces AP conduction in fibers that are not sensitive to Na_V_1.7 inhibition.

**Figure 4:**
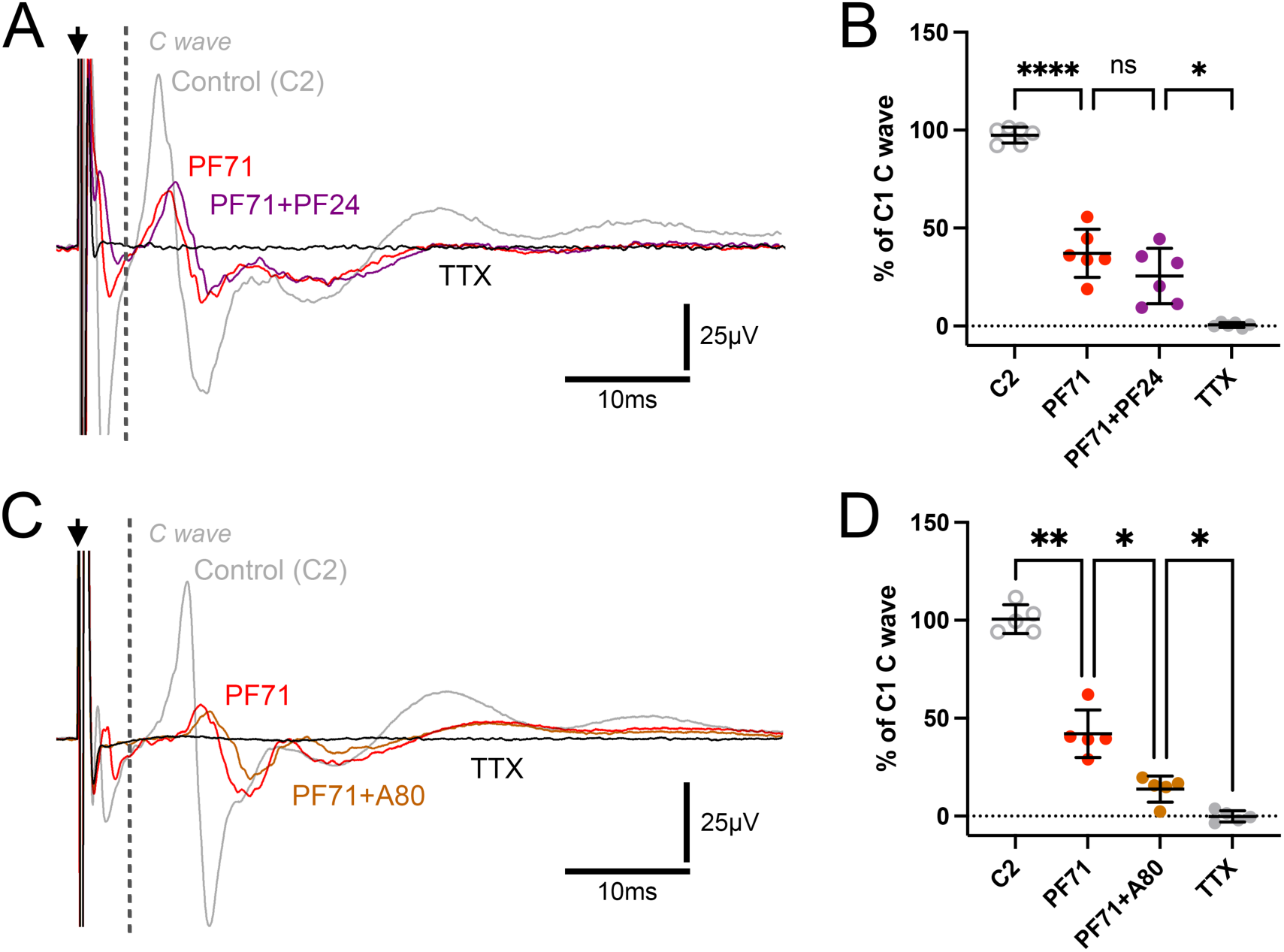
Effect of combination treatment with Na_V_1.7 inhibitor PF71 and Na_V_1.8 inhibitors on wild-type mouse vagal compound action potential (CAP). A, representative wild-type mouse vagal CAP during control C2 (grey trace), after 30mins treatment with PF71 (3µM, red trace), after PF71 (3µM) + PF24 (10µM) combination (purple trace), and 300nM TTX (black trace). B, mean ± SD of the AUC of the C-wave CAP normalized to the initial control (C1) during C2 control, PF71, combination PF71+PF24 and TTX (n=6). C, representative wild-type mouse vagal CAP during control C2 (grey trace), after 30mins treatment with 3uM PF71 (red trace), after PF71 (3µM) + A80 (1µM) combination (ochre trace), and 300nM TTX (black trace). D, mean ± SD of the AUC of the C-wave CAP normalized to the initial control (C1) during C2 control, PF71, combination PF71+A80 and TTX (n=5). In A and C, black arrow denotes the shock artifact; vertical grey line represents the division of A- and C-wave. In B and D, one-way repeated-measures ANOVA with Sidak’s multiple comparisons test *P<0.05, **P<0.005, ****P<0.0001, ns P>0.05.

### Action potential conduction in human vagal C-fibers

18 nerve samples from 18 different individuals (27.8% female) were obtained for our vagal CAP experiments. The median age of the patients was 53 years old (ranging from 6-79). Patient medical history and other relevant information are summarized in Table 1.

**Table 1:** Donor characteristics and general medical history.

| Age | Sex | Cause of death | Previous medical history |
|---|---|---|---|
| 6 | F | CV | Dandy Walker syndrome, Hypoxic-Ischemic Encephalopathy, gastrojejunostomy, tracheostomy, neurogenic bowel |
| 44 | M | CV | Nothing remarkable |
| 50 | M | Anoxia - seizure | adult-onset seizures, type 2 diabetes mellitus, hyperthyroidism |
| 40 | M | CV | Smoker, substantial alcohol, hypertension |
| 59 | M | CV | Ex-smoker, substantial alcohol |
| 56 | F | Pulmonary | Ex-smoker, hypertension, asthma/COPD, GERD, anemia, osteoarthritis |
| 57 | F | CV | Hypertension, high cholesterol, aortic regurgitation, colon cancer (remission), migraines, parathyroidectomy |
| 39 | M | CV | Type 1 diabetes mellitus, hypertension, high cholesterol, chronic kidney disease |
| 59 | M | Head trauma | High cholesterol, type 2 diabetes mellitus, hypothyroid, depression |
| 30 | M | Anoxia | Smoker, methamphetamine, cocaine, psychiatric hospitalization |
| 19 | M | Head trauma | Smoker |
| 63 | M | CV | Hypertension, high cholesterol, type 2 diabetes mellitus, depression |
| 71 | M | CV | Smoker, asthma, COPD, hypertension, high cholesterol, MI |
| 54 | M | CV | Smoker, cocaine, COPD, esophageal reflux |
| 53 | M | CV | Smoker, hypertension, congestive heart failure, stroke |
| 79 | F | CV | High cholesterol, hypothyroidism, breast cancer (remission) |
| 53 | M | CV | Smoker, methamphetamine, cocaine, stomach cancer (remission), hypertension, MI |
| 49 | F | CV | Smoker, COPD, hypertension |

Similar to mouse vagus nerves, 300nM TTX completely abolished the vagal C-fiber CAP signal (P=0.0008, Fig. 5A & 5B), consistent with a previous study conducted in human bronchial branches of the vagus nerve (Kollarik et al., 2022). The C-fiber CAP signal after TTX treatment was not significantly different from the CAP signal that was measured after the pinching of the nerve (TTX 300nM: 3.07±7.3% of C1 vs Pinch: 0% of C1, n=4, P=0.64). This once again indicates that TTX-S Na_V_1s are essential for all human vagal C-fiber AP conduction.

**Figure 5:**
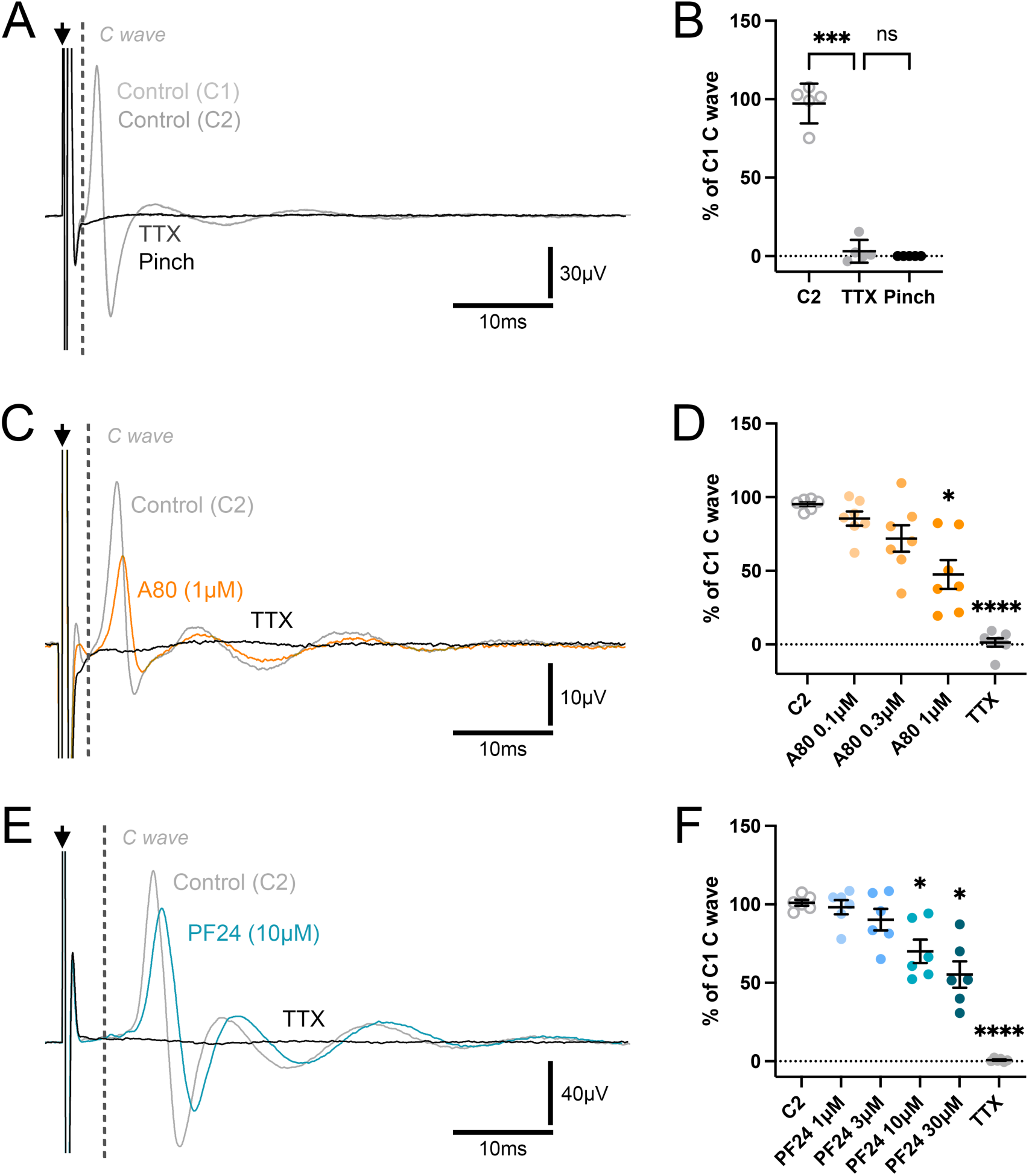
Effect of TTX and selective-Na_V_1.8 inhibitors on human vagal compound action potential (CAP). A, representative human vagal CAP during control C1 (light grey trace), control C2 (grey trace) and after 300nM TTX (dark grey trace) and nerve pinch (black trace). B, mean ± SD of the AUC of the C-wave CAP normalized to the initial control (C1) during C2 control, after TTX (300nM), and after nerve pinch (n=5). C: Representative human vagal CAP during control C2 (grey trace), after A80 (1µM, orange trace) and 300nM TTX (black trace). D, mean ± SD of the AUC of the C-wave CAP normalized to the initial control (C1) during C2 control, A80 (0.1 to 1µM) and TTX (n=7). E: Representative human vagal CAP during control C2 (grey trace), after 10µM PF24 (teal trace) and 300nM TTX (black trace). F, mean ± SD of the AUC of the C-wave CAP normalized to the initial control (C1) during C2 control, PF24 (1 to 30µM) and TTX (n=6). In A, C and E, black arrow denotes the shock artifact; vertical grey line represents the division of A- and C-wave. In B, D and F, one-way repeated-measures ANOVA with Sidak’s multiple comparisons test, compared to respective C2 control values *P<0.05, **P<0.005, ****P<0.0001, ns P>0.05.

Next, we tested whether TTX-R Na_V_1.8 was required for human C-fiber AP conduction as seen in mice. Concentration-inhibition curves for both A80 and PF24 were generated using human vagus nerve CAP recordings. Both drugs showed a concentration-dependent inhibitory effect on the C-fiber CAP signal (Fig. 5C–5F). 30-minute incubation with 1 µM A80 significantly reduced the human C-wave CAP (C2: 95.2±3.9% of C1 vs A80 1µM: 47.5±25.8% of C1, Fig. 5C & 5D, n=7, P=0.03). An unpaired comparison of the C-wave reduction by 1 µM A80 in human and mouse CAP experiments showed a trend for greater inhibition in human vagus nerves, although this did not reach statistical significance (Supplemental Fig. 3A, P=0.010). 30-minute incubation with 10 µM PF24 significantly reduced the human C-wave CAP (C2: 100.9±4.6% of C1 vs PF24 10µM: 70.1±18.2% of C1, Fig. 5E & 5F, n=6, P=0.028), with 30 µM PF24 inducing further inhibition. An unpaired comparison of the C-wave reduction by 10 µM PF24 in human and mouse CAP experiments showed similar levels of inhibition (Supplemental Fig. 3B, P=0.27). Thus, our findings demonstrate that the unexpected role of TTX-R Na_V_1.8 in conducting AP in mouse vagal fibers is conserved in human vagal C-fibers.

## Discussion

Na_V_1s are required for AP electrogenesis, facilitating both the initiation and conduction of APs (Catterall et al., 2005). AP initiation at peripheral terminals of sensory nerves occurs when a stimulus depolarizes the neuronal membrane past the Na_V_1 threshold potential (Mazzone & Undem, 2016). This depolarization opens Na_V_1s at the spike initiation zone, triggering Na+ influx, which results in the upstroke of the AP. The AP is then conducted along the nerve axon by sequential opening of neighboring axonal Na_V_1s. We have previously shown that vagal AP conduction is dependent on multiple voltage-gated sodium channels (Nair et al., 2023). Our data here provides evidence for the hypothesis that TTX-R Na_V_1.8 is required for AP propagation along a subset of C-fibers in both mouse and human vagus nerves. We further demonstrate that Na_V_1.8 is required for AP conduction in a subset of vagal nerve fibers that also require functional Na_V_1.7 channels.

Excessive or aberrant activity in vagus nerves has been implicated in many diseases affecting visceral organs such as asthma and gut dysmotility (Mazzone & Undem, 2016; Prescott & Liberles, 2022). Although the vagus nerve is a mixed peripheral nerve that contains both fast-conducting A-fibers and slow-conducting C-fibers, sensory C-fibers typically carry nociceptive information critical for mediating defensive reflexes. The C-wave component of a single vagal CAP is a summation of all the single APs conducted along each C-fiber within the vagus nerve. Action potentials are all-or-nothing events, thus a decrease in C-wave AUC represents a decrease in the number of C-fiber axons that have successfully propagated an AP. Na_V_1 inhibition in nociceptive nerves represents an attractive therapeutic strategy for treating diseases affecting vagally-innervated organs because it is theoretically independent of the triggering noxious stimulus. We have restricted quantification to the C-wave because A-wave signals often deteriorated during longer CAP experiments, particularly in human samples. It is possible that the surgical extraction and transportation might have led to the accelerated deterioration of the health of A-fibers as they are known to be more susceptible to hypoxic conditions (Clark et al., 1935; Jain et al., 2013).

Interpretation of these studies depends on the selectivity of the Na_V_1.8 inhibitors used. The published IC50s of Na_V_1 inhibitors are most often established using patch clamp experiments, where conditions such as extracellular sodium concentrations and resting membrane potentials are often manipulated to “push” the channels into particular gating states. Apart from TTX, which is a pore blocker (Shen et al., 2018), all the inhibitors used in this study exhibit state-dependent inhibition (Alexandrou et al., 2016; Jarvis et al., 2007; Payne et al., 2015). Hence, it is not certain that the reported selectivity of these inhibitors will be maintained in neuronal tissue under native conditions. Nevertheless, the concentrations we have used in this study have been tested in previous in vivo and ex vivo studies (Kollarik et al., 2018, 2022; Miranda-Morales et al., 2010; Nair et al., 2023).

TTX has been classically used to study the role of Na_V_1s in AP initiation and conduction. Multiple studies have demonstrated that TTX completely abolished AP conduction in all vagal afferents innervating the mouse esophagus (Hadley et al., 2021), the guinea pig esophagus (Ru et al., 2020), the guinea pig lungs (Muroi et al., 2011) and the human bronchi (Kollarik et al., 2022), as well as in all vagal efferents innervating the mouse, guinea pig and human bronchi (Kocmalova et al., 2017) and the guinea pig esophagus (Pavelkova et al., 2020). Such findings led to the conclusion that only TTX-S Na_V_1s were involved in AP conduction in vagal nerve fibers. Our experiments here corroborated those findings as TTX again abolished electrically evoked AP conduction in both mice and human vagus nerves. However, the TTX-S Na_V_1 requirement does not preclude a necessary role for Na_V_1.8 in AP conduction (Nair et al., 2023). Here we found that Na_V_1.8 inhibitor PF24 caused ∼40% reduction in electrically stimulated C-fiber AP conduction in mouse vagal nerves. Since PF24 did not have any effect on C-fiber AP conduction in our Na_V_1.8 KO mice experiments, we conclude that Na_V_1.8 is required to conduct APs by a subset of mouse vagal nerves.

NaV1 isoform expression in vagal afferent neurons is heterogeneous. Almost all vagal neurons express Na_V_1.7 and Na_V_1.8, but considerable populations also express Na_V_1.1, 1.2, 1.3, 1.6 and 1.9 (Hadley et al., 2021; Kollarik et al., 2018; Kupari et al., 2019; Kwong et al., 2008). Na_V_1.7 is critical for AP conduction in ∼66% of vagal nerve fibers (Kollarik et al., 2022; Nair et al., 2023). The remaining TTX-S AP conduction in the mouse vagus was inhibited in our previous study by ICA (an inhibitor of Na_V_1.1, Na_V_1.3 and NaV1.6 in mice) (Nair et al., 2023). Here, following Na_V_1.7 inhibition with PF71, the addition of PF24 evoked no further reduction in the C wave CAP, indicating that Na_V_1.8 is essential for AP conduction in only those vagal fibers that express and employ Na_V_1.7 for AP conduction.

While unexpected, the requirement for Na_V_1.8 in AP conduction in a subset of vagal nerve fibers could be explained by its well-documented kinetics and biophysical properties during AP electrogenesis (Catterall et al., 2005). The ability of Na_V_1.8 to support AP initiation independently was demonstrated in studies of airway vagal C-fibers projected from jugular neurons, where AP initiation is resistant to TTX and blocked by PF24 (Kollarik et al., 2018). Moreover, a functional role for axonal Na_V_1.8 has been described in TTX-resistant axons in rodent sciatic and spinal nerves (Gold et al., 2003; Klein et al., 2017; Palomino et al., 2026; Pinto et al., 2008). Mechanistic patch-clamp studies in nociceptor DRG neurons of rodents have demonstrated that Na_V_1.7 is open at more hyperpolarized potentials to set the AP threshold (Blair & Bean, 2002; Rush et al., 2007). In contrast, Na_V_1.8, opening at more depolarized membrane potentials, is responsible for most of the inward current during AP upstroke (Blair & Bean, 2002; Matsutomi et al., 2006; Renganathan et al., 2001). In the context of vagal AP conduction, we speculate the existence of a population of fibers that employ both Na_V_1.7 and Na_V_1.8 for the depolarization needed for conducting APs. Blocking Na_V_1.8 most likely stunts the required inward current needed for the depolarization of the neighboring axonal segment. In the same fibers, application of TTX abolishes AP conduction because they cannot reach AP threshold without Na_V_1.7. Fibers that maintain AP conduction in the presence of Na_V_1.8 inhibition likely have a high enough density of Na_V_1.7 and/or other TTX-S isoforms. Alternatively, such fibers may employ post-translational modifications or Na_V_ β isoforms to alter Na_V_1 TTX-S isoform gating properties and eliminate the Na_V_1.8 requirement for AP conduction (Barry et al., 2014; Loya-López et al., 2022; Powell et al., 2026; Shirahata et al., 2006; Vijayaragavan et al., 2004).

Here A80 also inhibited the vagal AP conduction in mouse vagus nerves. However, some inhibition was also observed in Na_V_1.8 KO mouse vagus nerves. Together, these data suggest A80 blocked Na_V_1.8 and non-Na_V_1.8 channels to achieve its AP conduction inhibition in the vagus nerve. Even though A80 is considered selective for Na_V_1.8 (>100-fold selective for hNa_V_1.8 vs hTTX-S channels), there have been reports of its potential off-target effects in vivo and ex vivo. In thin fiber somatic afferents (also known as group III and IV afferents) innervating muscle spindles that do not express Na_V_1.8, A80 significantly reduced pressor reflexes to capsaicin and lactic acid (Stone et al., 2013). This group postulated that Na_V_1.6, which is expressed by these muscle afferents could be a potential target for A80 at the administered dose. Due to the close evolutionary lineage shared by Na_V_1.8 and Na_V_1.5, A80 has also been shown to have an effect on cardiomyocytes and HEK cell line expressing Na_V_1.5 (Han et al., 2015). Although Na_V_1.5 is often considered a non-neuronal, cardiac-only TTX-R Na_V_1, some vagal sensory neurons express this isoform in transcriptomic analyses (Bai et al., 2019; Kupari et al., 2019).

The phenomenon where both TTX-S Na_V_1s and Na_V_1.8 are required for AP conduction in the same axons was observed in human vagus nerve experiments as well. Despite the observation that human vagal AP conduction was abolished by TTX, consistent with previous studies (Kollarik et al., 2022), both A80 and PF24 demonstrated significant inhibitory effect on electrically-evoked C-fiber AP conduction. In previous patch clamp experiments, PF24’s affinity for human TTX-R was only ∼1.67-fold higher than that for mouse TTX-R channels (IC50s: ∼0.3nM vs ∼0.5nM respectively) (Payne et al., 2015). Here we found little difference in the PF24 inhibition of vagal AP conduction in mouse and human nerves. Although the patch-clamp profile of A80 on mouse Na_V_1.8 channels is unavailable, data exists for recombinant rat Na_V_1.8 channels (Jarvis et al., 2007), which share 96% homology with the mouse ortholog. Comparing the IC50s of A80 against human Na_V_1.8 with rat Na_V_1.8 in their inactivated states, A80 demonstrated ∼6-fold higher affinity towards human Na_V_1.8 (89% homology with rat ortholog). In our recordings, there was a trend towards greater A80-evoked inhibition in human vagus nerves compared to the mouse, although this difference did not reach statistical significance (P=0.1). This could reflect A80’s higher affinity for human Na_V_1.8 or it could be due to A80’s inhibition of other Na_V_1s. However, it is important to note that we have provided no evidence of A80’s selectivity for human non-Na_V_1.8 isoforms. Furthermore, the role of non-Na_V_1.7 TTX-S isoforms in AP conduction in human nerves has not been directly studied.

The clinical importance of vagal afferent Na_V_1.8 is supported by a study in guinea pigs, that demonstrated the inhalation of A80 caused a significant reduction in the number of coughs produced in response to capsaicin (Brozmanova et al., 2019). Interestingly, systemic administration of A80 caused a greater inhibition of capsaicin-induced coughs compared to inhaled A80. Therefore, it is possible that systemic A80 could have targeted axonal Na_V_1.8 expressed on airway vagal fibers, in addition to the nerve terminals in the large airways and lungs accessed by the inhaled inhibitor. Nevertheless, clinical studies of patients with loss-of-function Na_V_1.8 mutations do not report any gross autonomic dysregulation (Dabby et al., 2016; Huang et al., 2013; Kist et al., 2016) which could be expected with ∼35% reduction in the number of vagal fibers capable of conducting APs. A possible explanation is that congenital loss-of-function mutations affecting Na_V_1 isoforms may trigger compensatory changes in the expression of other Na_V_1 isoforms, which can subsequently assume the compromised isoform’s function. Such compensatory changes have been reported in DRG neurons of the Na_V_1.8 KO mouse (Akopian et al., 1999). Suzetrigine, an FDA-approved Na_V_1.8-selective inhibitor, has also not been associated with obvious autonomic adverse effects in clinical trials. While our experiments only used desheathed vagus nerves, in vivo peripheral nerves are ensheathed with perineurium and epineurium. It is well known that such nerve axon barriers hinder access of drugs (Hackel et al., 2012). One possible reason for the lack of autonomic side effects is that these barriers prevent Suzetrigine from reaching the required concentration at axonal sites needed to impair vagal AP conduction.

In summary, these studies use two distinct Na_V_1.8-selective inhibitors to show that axonal Na_V_1.8 is essential for AP conduction in a subset of C-fibers that are also sensitive to Na_V_1.7 inhibition. While the therapeutic targeting of AP conduction in the vagus nerve per se may lack the specificity of targeting AP initiation in afferent terminals innervating specific organs, it is likely that effective blockade of either isoform within the vagus nerve would likely have widespread effects on multiple integrative autonomic systems. In addition, Na_V_1.8 expression has been shown to be upregulated in injured sciatic and infraorbital nerves (Mulpuri et al., 2022; Ruangsri et al., 2011; Thakor et al., 2009), and it is possible that inflammatory disease or injury may further increase the role of axonal Na_V_1.8 expression in vagal AP conduction.

## Author contributions

TET-C conceived the project; TET-C and SSN designed the research; SSN performed the experiments; SSN and TET-C analyzed the data; SSN and TET-C prepared the figures; SSN drafted the manuscript; SSN and TET-C edited and revised the manuscript; SSN and TET-C approved the final version.

