## Supplemental Figures for "Tetrodotoxin-resistant Na_V_1.8 contributes to action potential conduction in a population of mouse and human vagal nerve fibers"

**Supplemental figures with legends**

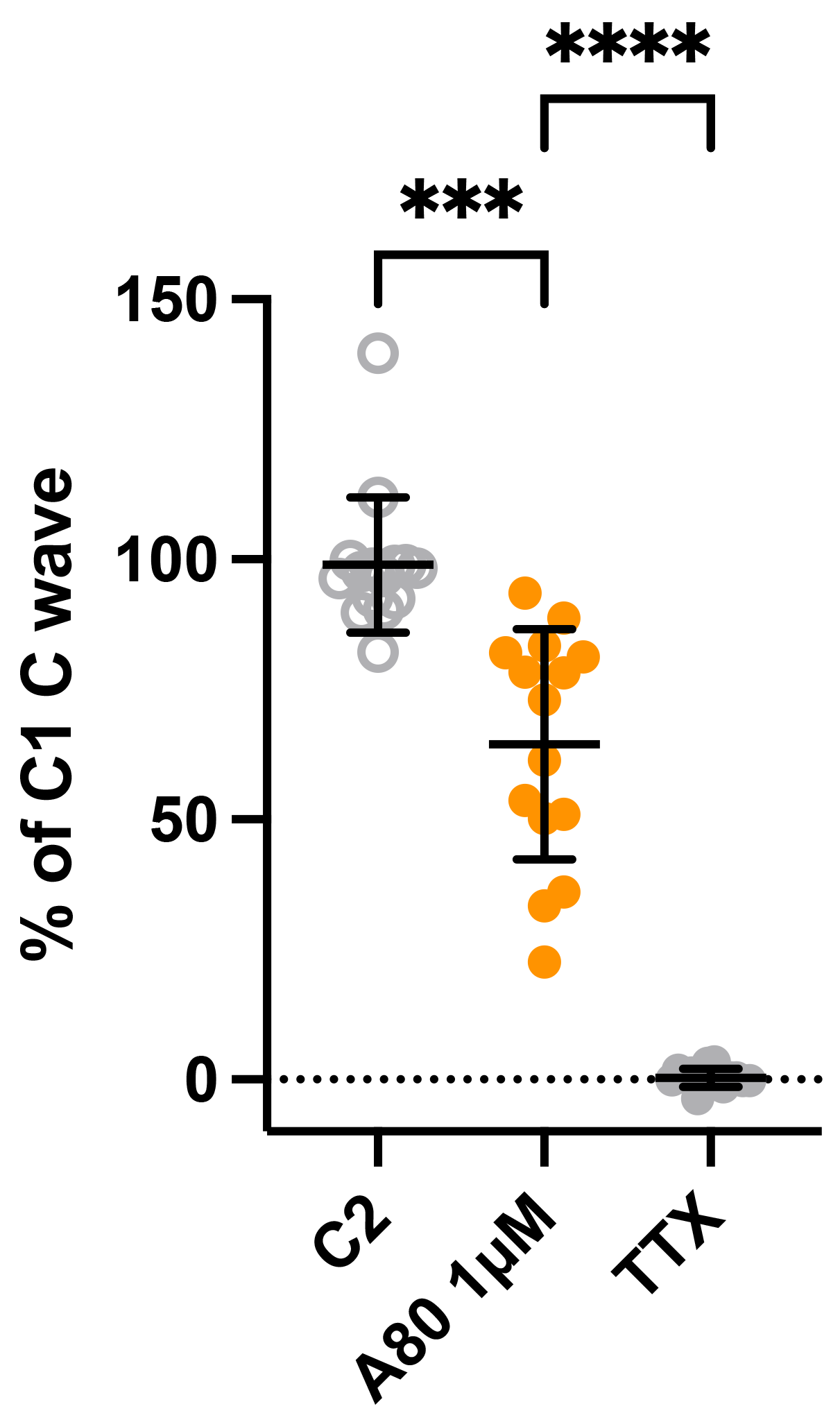

Figure S1: Effect of 1µM A80 on wild-type mouse vagal compound action potential (CAP). Mean ± SD of the AUC of the C-wave CAP normalized to the initial control (C1) during C2 control, 1µM A80 and 300nM TTX (n=15). One-way repeated-measures ANOVA with Sidak’s multiple comparisons test ***P<0.0005, ****P<0.0001

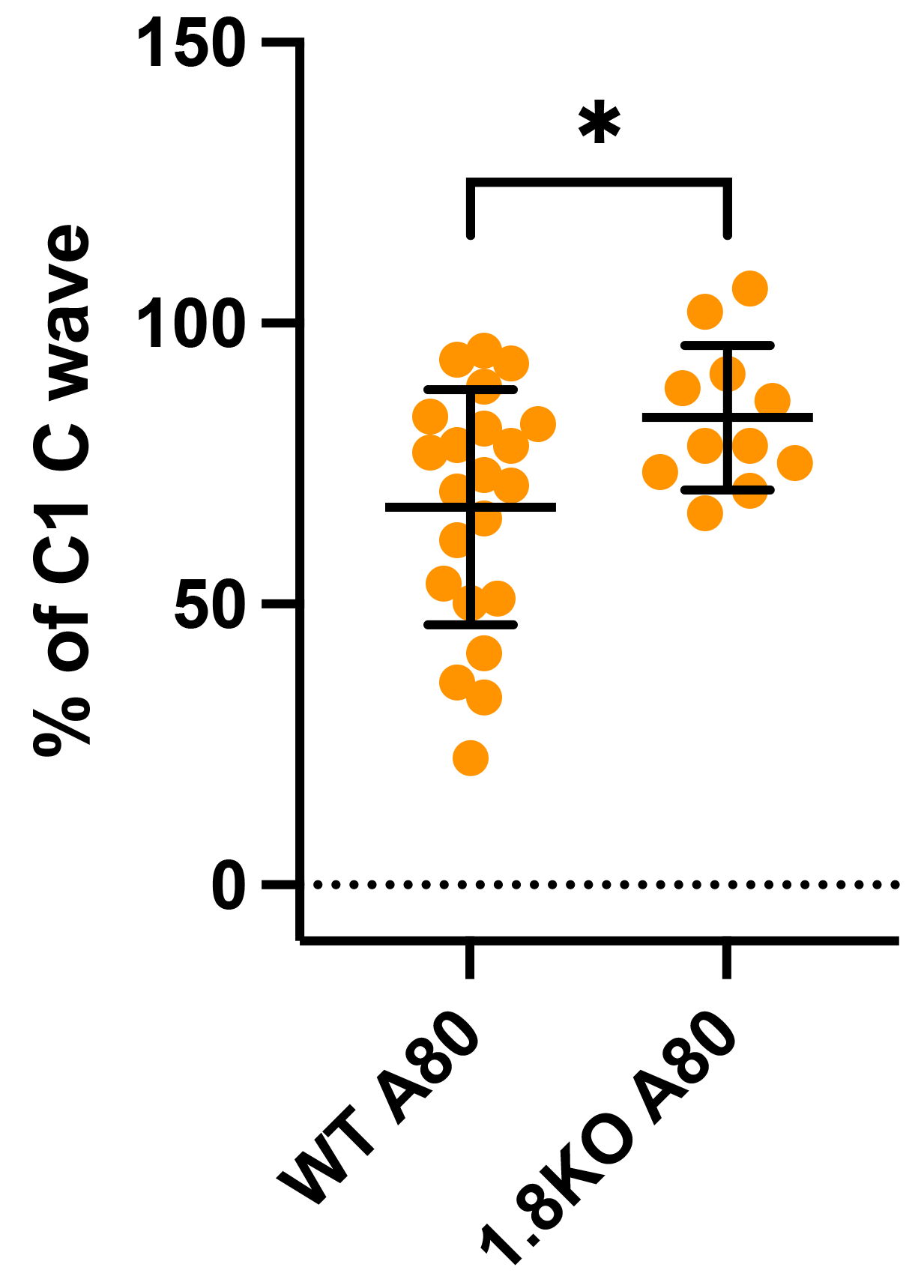

Figure S2: Comparison of the inhibitory effect of 1µM A80 inhibitory on wild-type and Na_V_1.8 KO mouse vagal compound action potential (CAP). Mean ± SD of the AUC of the C-wave CAP normalized to the initial control (C1). Data are pooled from multiple experiments: Wild-type n=22, Na_V_1.8 KO n=11. Unpaired t-test with Welch’s correction *P<0.05.

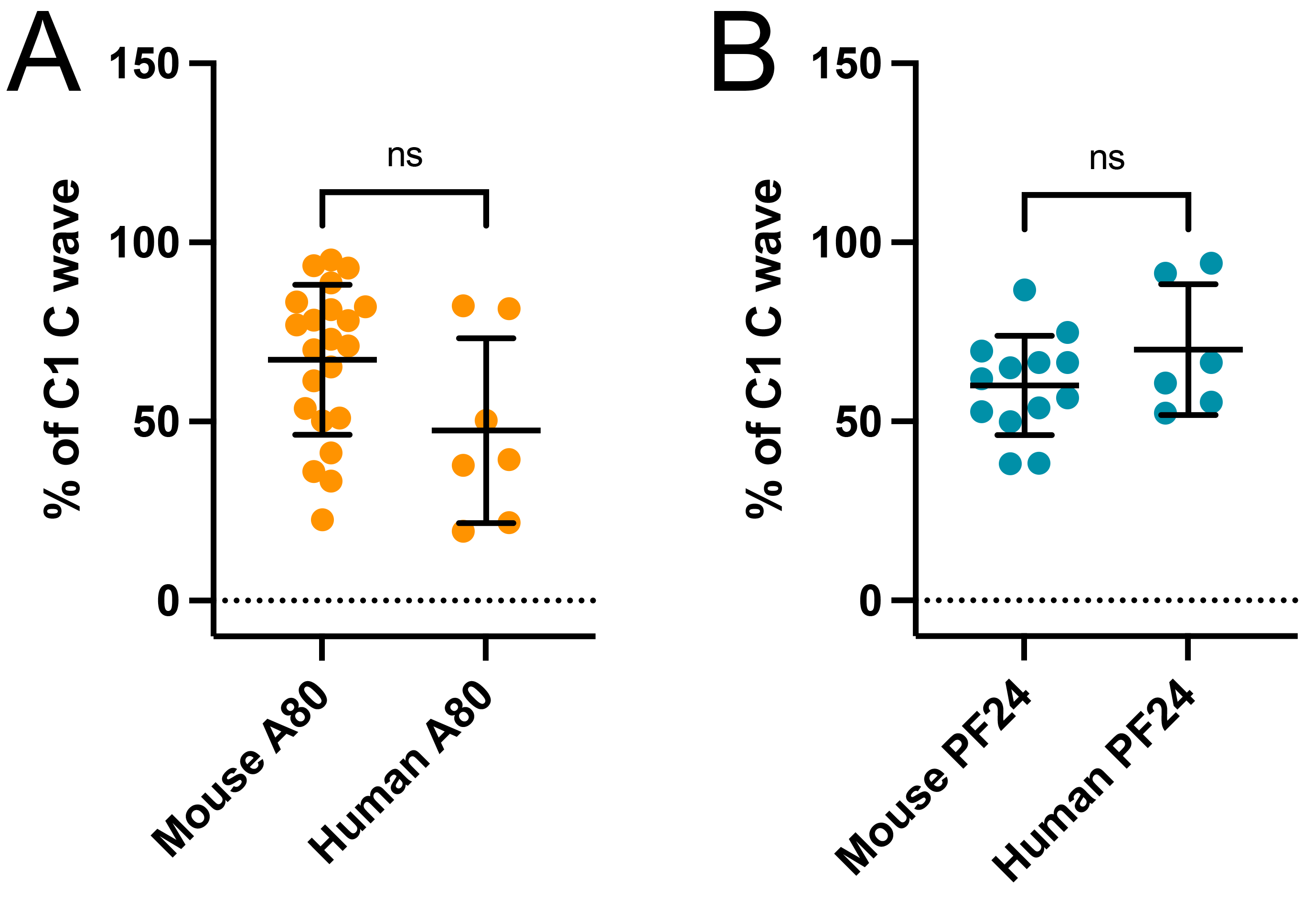

Figure S3: Inter-species comparison of the effect of Na_V_1.8 inhibitors on mouse and human vagal compound action potential (CAP). Data are pooled from multiple experiments. A, mean ± SD of the AUC of the C-wave CAP normalized to the initial control (C1) during 1µM A80 on mouse (n=22) and human vagal CAP (n=7). B, mean ± SD of the AUC of the C-wave CAP normalized to the initial control (C1) during 10µM PF24 on mouse (n=13) and human vagal CAP (n=6). Unpaired t-test with Welch’s correction, ns P>0.05
